# Memory load influences our preparedness to act on visual representations in working memory without affecting their accessibility

**DOI:** 10.1101/2024.02.23.581707

**Authors:** Rose Nasrawi, Mika Mautner-Rohde, Freek van Ede

## Abstract

It is well established that when we hold more content in working memory, we are slower to act upon specific content when it becomes relevant for behavior. Here, we asked whether slower responses with higher working-memory load can be accounted for by slower access to sensory representations held in working memory (according to a serial internal search), or by a lower preparedness to act upon them. To address this, we designed a visual-motor working-memory task in which participants memorized the orientation of two or four colored bars, of which one was cued for reproduction. We independently tracked the selection of visual (cued item location) and motor (relevant manual action) information from the EEG time-frequency signal, and compared their latencies between our memory-load conditions. We confirm slower memory-guided behavior with higher working-memory load and show that this is associated with delayed motor selection, but find no evidence for a concomitant delay in the latency of visual selection. Moreover, we show that variability in decision times within each memory-load condition is associated with corresponding changes in the latency of motor, but not visual selection. These results reveal how memory load affects our preparedness to act on, without necessarily changing the ability to access, sensory representations in working memory. This posits action readiness is a key factor that shapes the speed of memory-guided behavior and that underlies delayed responses with higher working-memory load.

## INTRODUCTION

Working memory is a fundamental cognitive process that enables us to retain relevant sensory information in order to guide upcoming behavior (Baddeley, 1992; Chatham & Badre, 2015; D’Esposito & Postle, 2015; Myers et al., 2017; van Ede & Nobre, 2023). To delineate the foundational mechanism of working memory, working-memory load has frequently been experimentally manipulated – i.e., the number of memory items to be retained by participants (e.g., Alvarez & Cavanagh, 2004; Awh et al., 2007; Bays & Husain, 2008; Luck & Vogel, 1997, 2013; Todd & Marois, 2004; Xu et al., 2018). Among others, these studies have consistently reported that an increase in working-memory load is associated with poorer memory-guided behavior for individual memory items.

Specifically, when we hold more content in working memory, we are slower to act upon specific content when it becomes relevant for behavior. One classic example of this comes from the Sternberg task (Sternberg, 1966a, 1969, 1975). In this task, participants sequentially memorized a set of items and determined whether a target item was part of the original set. This revealed that, as more items were held in working memory, participants were linearly slower to respond. These findings have typically been interpreted to reflect a serial internal-comparison process (Sternberg, 1966; see also: Kong & Fougnie, 2019): items in working memory are searched one-by-one, and as a consequence, this process takes longer when there are more items to search among. From this perspective, it follows that slower responses with an increase in memory load go along with *slower access to the sensory representations* held in visual working memory.

Simultaneously, it is essential to recognize that working memory is not merely a storage for sensory information, but also an anticipatory buffer that forms the bridge between perception and action (Chatham & Badre, 2015; Formica et al., 2024; Heuer et al., 2020; Myers et al., 2017; Olivers & Roelfsema, 2020; van Ede, 2020; van Ede & Nobre, 2023). Accordingly, the retention of visual information in working memory often takes place alongside planning for upcoming behavior. For example, a recent series of studies utilizing a novel working-memory task in which sensory objects were linked to prospective actions (van Ede et al., 2019), demonstrated that, alongside the encoding and retention of visual information, action planning occurs during working memory and correlates with decision times (Boettcher et al., 2021; Nasrawi et al., 2023; Nasrawi & van Ede, 2022). As such, a viable complementary account for delayed responses with higher working-memory load is that: when we hold more items in working memory, it is harder to prepare for action. From this alternative perspective, slower responses with an increase in memory load may equally well be driven by a decreased *preparedness to act* on content in working memory, irrespective of the time it takes to access this content.

To disentangle these alternative scenarios – item accessibility versus action readiness – we used electroencephalography (EEG) to track the temporal dynamics of visual and motor selection, serving as markers for accessibility and readiness, respectively. Previewing our results, we first confirm robust delayed responses with higher working-memory load (four versus two items). Strikingly, with higher working-memory load, we found no evidence for a change in the latency of visual selection (item location), while we observed clear evidence for a delayed selection of motor information (required response hand). Moreover, within each memory load condition, we observed that variability in the onset of memory guided behavior (fast/slow) was associated with corresponding changes in the latency of motor, but not visual selection. These neural dynamics provide direct evidence that working-memory load can affect our preparedness to act on, without necessarily changing our ability to access, sensory representations in working memory.

## RESULTS

Participants performed a visual-motor working-memory task (**Figure 1**; building on Boettcher et al., 2021; Nasrawi et al., 2023; van Ede et al., 2019) in which they were asked to memorize the orientation of two or four colored bars. At a given trial (**Figure 1A**), four colored bars were shown at encoding, of which one was cued for reproduction after a working-memory delay, indicated by a change in the color of the central fixation cross. Participants used either of two buttons to turn the handles of a response dial clockwise or anti-clockwise to precisely reproduce the orientation of the cued item. We manipulated working-memory load across blocks by instructing participants to selectively attend and memorize either two items (load-two) or all four items (load-four), indicated by their colors (**Figure 1B**). Additionally, we manipulated the tilt-response mapping, so that a certain bar tilt required a reproduction report with a certain response hand (**Figure 1C**). This was achieved by a block-wise manipulation of the starting position of the response dial (vertical/horizontal). In blocks in which the dial started in the vertical position, reproducing a leftward tilt required a response with the left hand, and vice versa for a rightward tilt. Conversely, in blocks in which the dial started in the horizontal position, reproducing a leftward tilt required a response with the right hand, and vice versa for a leftward tilt. Consequently, within a given block, participants could plan ahead which response hand would be needed to reproduce a certain item’s tilt (while ensuring that across blocks tilt and response hand remained orthogonal, as in: Nasrawi et al., 2023).

**Figure 1.**
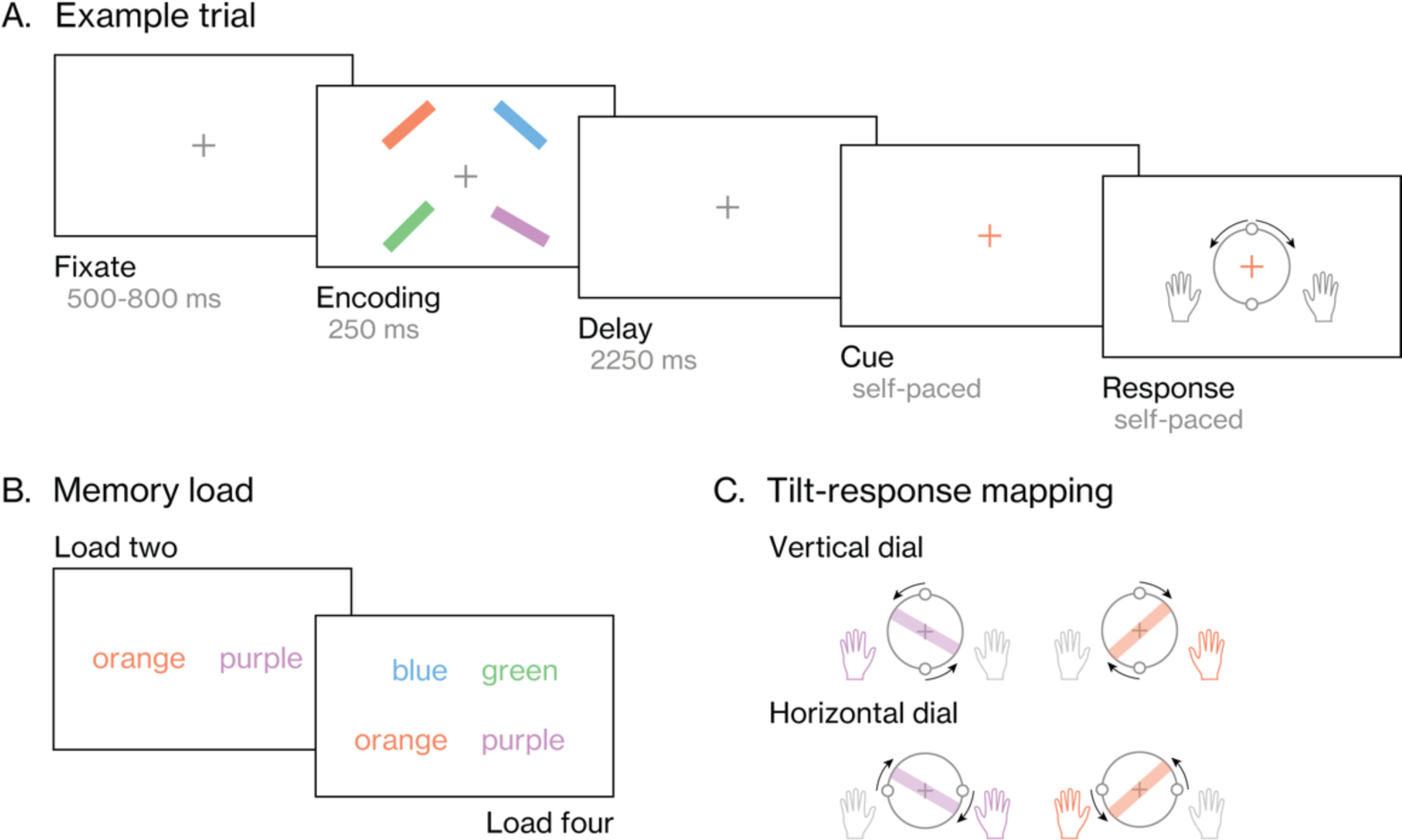
Visual-motor working memory task. **(A) Example trial.** Participants viewed four colored oriented bars, which they were instructed to memorize. After a brief delay, the fixation cross changed color, cueing participants to reproduce the orientation of the memorized bar that matches this color cue. Participants gave their response by turning the handles of a response dial. These handles could be turned clockwise by pressing and holding the ‘M’ key with their right hand, or counterclockwise by pressing and holding the ‘Z’ key with their left hand. **(B) Memory load manipulation.** Preceding a block of trials, participants were instructed to selectively memorize either two or four bars based on their color, of which in all cases only one would be cued for reproduction. **(C) Tilt-response mapping manipulation.** The starting position of the response dial was manipulated between blocks: it could be either vertical, or horizontal. Given that participants could only turn the dial handles a maximum of 90 degrees in either direction, the mapping between tilt and response hand varied throughout the experiment. Given a vertical response dial, a leftward tilted bar could only be reproduced using the left hand, and a rightward tilted bar using the right hand. Alternatively, given a horizontal response dial, a leftward tilted bar could only be reproduced using the right hand, and a rightward tilted bar using the left hand.

### Working-memory load affects task performance

Before turning to our main question regarding the dynamics of visual and motor selection as a function of working-memory load, we looked into the effect of working-memory load (two/four items) on decision times (ms) and absolute errors (degrees) of the tilt-reproduction reports. Decision time was defined at the time between cue onset (i.e., color change of the fixation cross) and participants’ response initiation. Absolute error was defined as the difference in orientation between the cued item and the reported orientation.

Participants were significantly slower to initiate their response in load-four trials, compared to load-two trials (**Figure 2A**; *t*(24) = 8.72, *p* < 0.001, Cohen’s *d_z_* = 1.74; see Lakens, 2013). Similarly, their orientation-reproduction reports were significantly less precise in load-four, compared to load-two (**Figure 2B**; *t*(24) = 8.06, *p* < 0.001, *d_z_* = 1.61). On average, participants were 197±24 (*M*±*SE*) ms slower and 5.92±0.67 (*M*±*SE*) degrees less precise in load-four compared to load-two. Additional visualization of these results showed how the distribution of decision times shifted to longer decisions times with load-four (**Figure 2A**, *lower panel*) and how participants reported the cued orientation in a fine-grained manner in both memory-load conditions, but with less overall precision in load-four (**Figure 2B**, *lower panel*). These results confirm that participants selectively memorized either two or four items, as instructed, despite the encoding display always containing four items.

**Figure 2.**
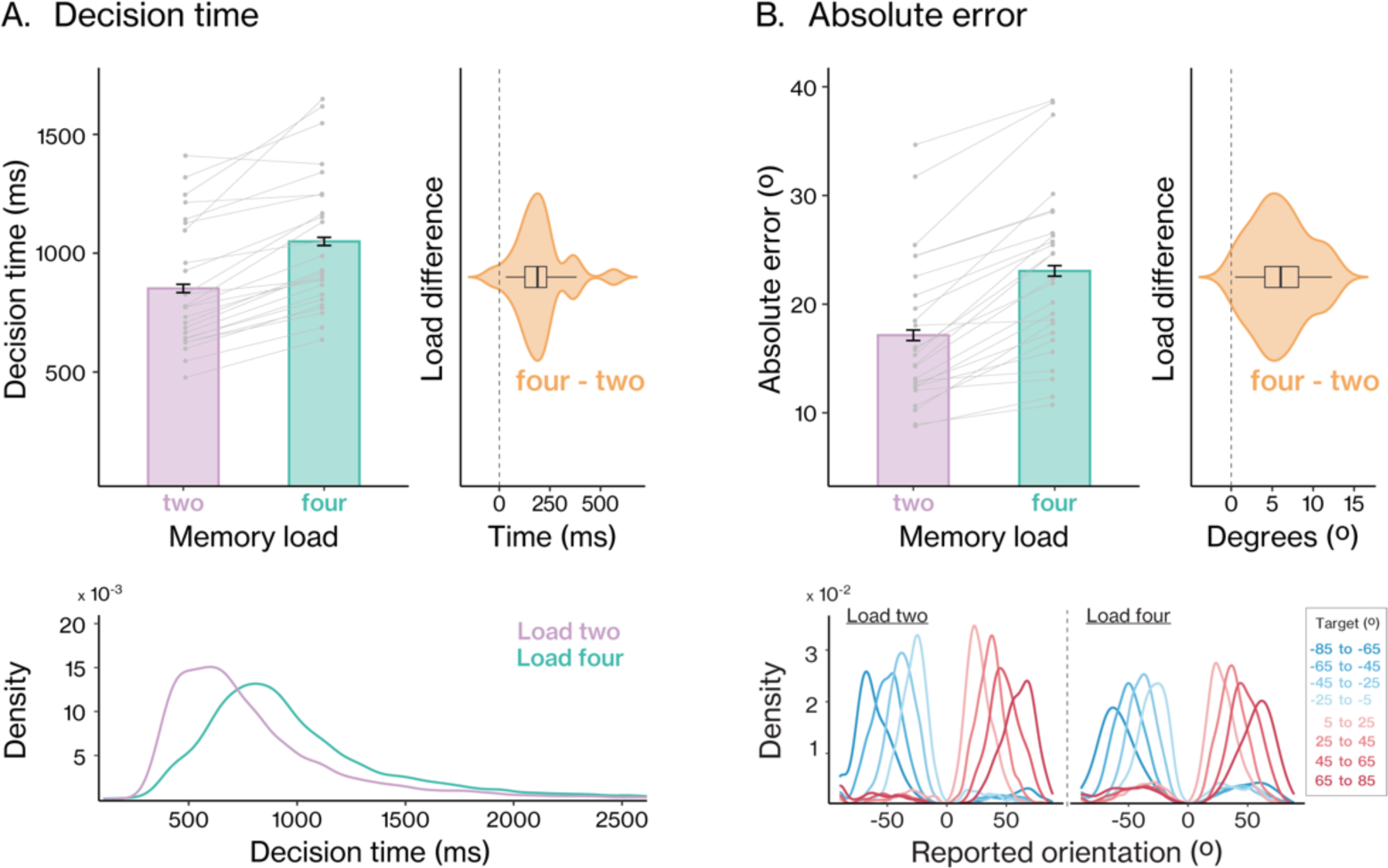
Memory load affects memory-guided behavior. **(A) Decision times.** Bar graphs show the average decision time (ms) as a function of memory load (two/four). Error bars represent within-subjects standard error (SE). Grey lines and points show individual participant data. The violin plot shows the effect of memory load (four-two). Density plots show the distribution of decision times in load-two and load-four. **(B) Absolute error**. Bar graphs show the average error (degrees) as a function of memory load (two/four), and violin plot shows the effect of memory load (four-two). Density plots show the distribution of reported-orientation (degrees) as a function of target orientation (colored, in 20-degree bins), for load-two and load-four.

While ample studies have targeted the reasons why performance gets worse with higher working-memory load (e.g., Bays et al., 2009; Bays & Husain, 2008; Ma et al., 2014; Oberauer et al., 2012; Oberauer & Lin, 2017; van den Berg et al., 2012), we were here particularly interested in what makes participants slower: does it take longer to access the relevant memory content (when there are more contents to select amongst) or is it simply harder to prepare to act on memory contents when there are more potential courses of action? To address this central question, we next turned to time-resolved signatures of visual and motor selection from working-memory.

### Selection of both visual and motor memory attributes after cue onset

Next, we looked into the dynamics of visual and motor selection after cue onset with EEG time-frequency analysis, both in load-two and load-four. An established neural marker of visual selection in scalp EEG measurements is the lateralization of alpha-band activity in posterior electrodes relative to the location of the selected item (e.g., Sauseng et al., 2005; Thut et al., 2006; Worden et al., 2000), including when selected items are held in visual working memory (Boettcher et al., 2021; Liu et al., 2022; Nasrawi et al., 2023a; Poch et al., 2017; van Ede et al., 2019; Wallis et al., 2015). Building on these prior studies, we here used posterior alpha lateralization as a marker of “visual selection”.

Similarly, an established marker of action selection in the EEG is the attenuation of beta-band activity in central electrodes contra-versus ipsilateral to the required response hand (e.g., Mcfarland et al., 2000; Neuper et al., 2006; Salmelin & Hari, 1994; Van Wijk et al., 2009) that has recently also been studied in the context of visual working memory (e.g., Boettcher et al., 2021; Ester & Weese, 2022; Nasrawi et al., 2023; Rösner et al., 2022; Schneider et al., 2017; van Ede et al., 2019). Building on these studies, we here used central beta lateralization as a marker of “motor selection”.

As shown in **Figure 3A**, we observed clear relative attenuation of alpha-band activity contra-versus ipsilateral to the cued items’ memorized location in electrodes PO7/PO8, shortly after cue onset, both in load-two (*upper panel*, cluster *p* = 0.0001) and in load-four (*lower panel*, cluster *p* = 0.0006). Topographies show the difference between trials where the cued item had occurred on the left versus right side of the display, and confirm that the attenuated alpha-band (8-12Hz) activity was particularly prominent in the left and right posterior electrodes, consistent with reflecting visual selection.

**Figure 3.**
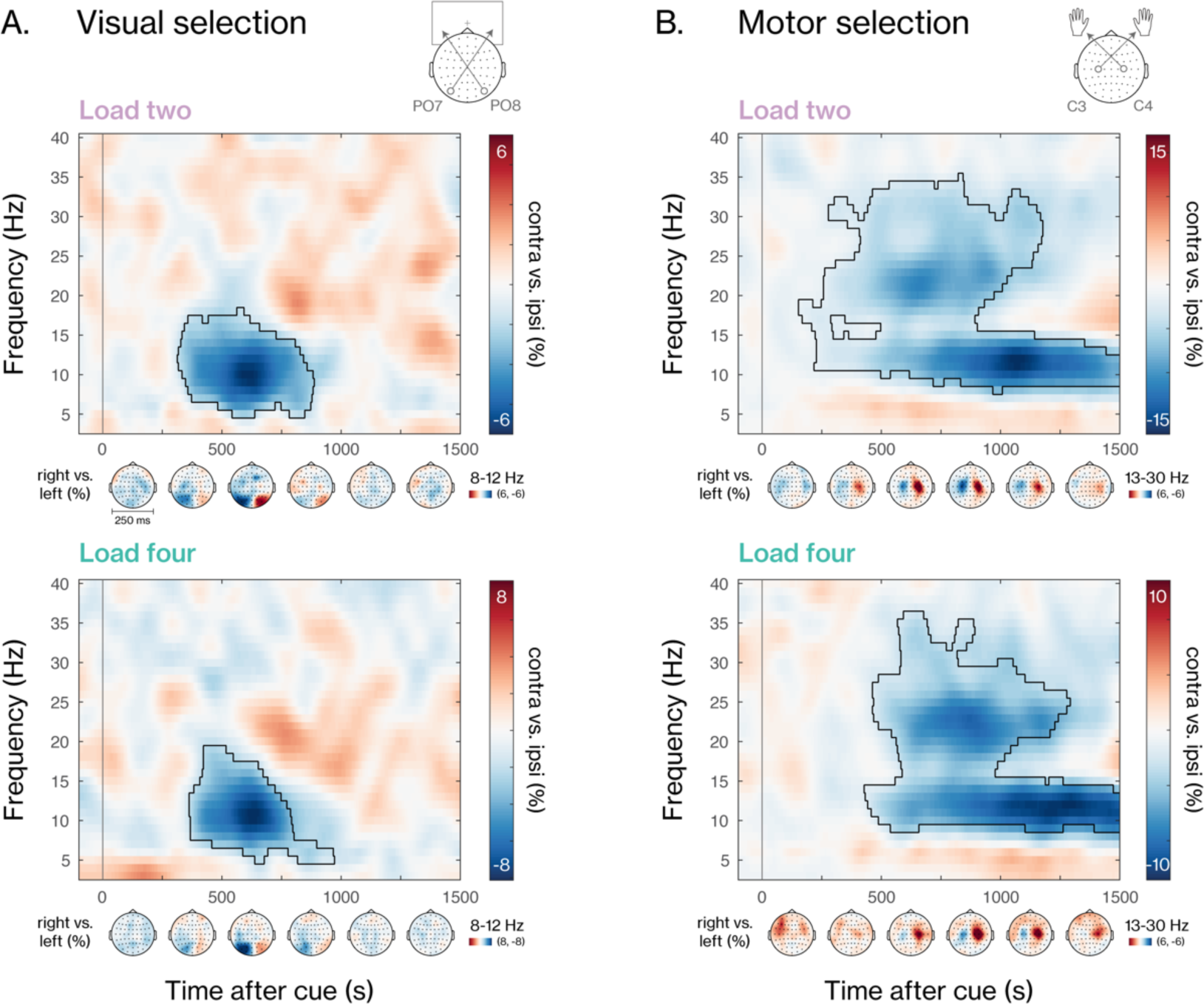
Selection of both visual and motor information after cue onset. **(A) Visual selection.** Lateralized time-frequency spectrum contra-vs ipsilateral to the cued items’ location (left/right) in electrodes PO7/PO8. Topographies show 8-12Hz lateralization (left versus right item location), averaged in steps of 250ms after cue onset. **(B) Motor selection.** Lateralized time-frequency spectrum contra-vs ipsilateral to the cued items’ required response hand (left/right hand) in electrodes C3/C4. Topographies show 13-30Hz lateralization (left versus right required response hand), averaged in steps of 250ms after cue onset. Black outlines indicate significant clusters yielded from cluster-based permutation. Black vertical lines indicate the onset of the cue (i.e., change in the color of the fixation cross). Upper panels show the data for load-two trials; lower panels show the data for load-four trials.

Similarly, after cue onset, we observed clear relative attenuation of beta-band activity (**Figure 3B**) contra-versus ipsilateral to the required response hand associated with the cued item in electrodes C3/C4, both in load-two (*upper panel*, cluster *p* = 0.0001) and in load-four (*lower panel*, cluster *p* = 0.0001). Topographies show the difference between trials that required a left versus right response hand, and confirm that the attenuated beta-band (13-30Hz) signatures were particularly prominent in the left and right central electrodes, consistent with reflecting action selection.

These data reveal robust neural markers of both visual-spatial attention to the cued item (visual selection), and selection of the required response hand (motor selection) in both load conditions. We next turn to our central question whether and how the respective temporal dynamics of visual and motor selection depend on the number of items held in working-memory.

### Memory load delays motor but not visual selection from working memory

Next, we aimed to answer our main question: are the observed slower onsets of the memory-guided behavioral reports with higher working-memory load (**Figure 2A**) driven by slower access to the appropriate (cued) sensory representations, or a lower preparedness to act on them?

To answer this question, we obtained time-courses of the averaged 8-12 Hz alpha-band lateralization in posterior electrodes (load-two cluster *p* < 0.0001; load-four cluster *p* = 0.002), and of the averaged 13-30 Hz beta-band lateralization in central electrodes (load-two cluster *p* = 0.0009; load-four cluster *p* = 0.0024). As shown in **Figure 4**, we found virtually indistinguishable time-courses of visual selection when overlaying selection dynamics between load-two and load-four (**Figure 4A**; no cluster found). In contrast, we did observe clear differences in motor selection, with later motor selection in load-four (**Figure 4B**; as can also be appreciated by revisiting the full time-frequency plots shown in **Figure 3)**. This was confirmed by a significant difference in motor selection between load-two and load-four trials (cluster p = 0.01).

**Figure 4.**
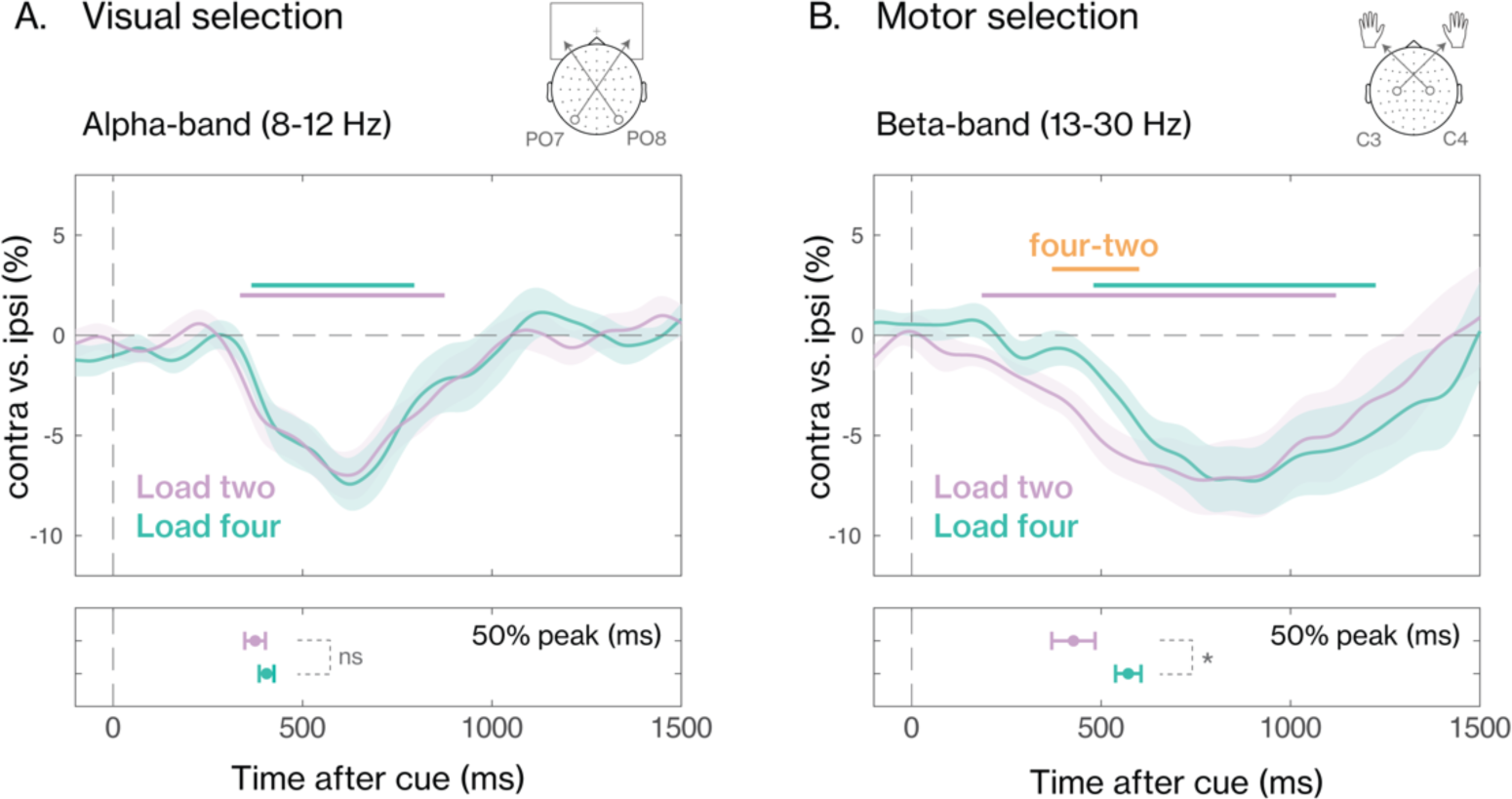
Memory load delays the selection of motor, but not visual information from working memory. Time-courses of **(A)** the lateralized visual alpha-band (8-12Hz) and **(B)** motor beta-band (13-30Hz) activity, for load-two (pink) and load-four (blue). The colored horizontal lines indicate significant clusters from the cluster-based permutation. The grey horizontal line shows the significant cluster for the difference between load-four and load-two. Black vertical dashed lines indicate the onset of the color cue. Shadings indicate standard error (SE) across participants. Lower panels show averaged jackknife estimates of the time the visual and motor time-courses first reach 50% of their minimum (or: negative peak). Error bars indicate standard errors.

To statistically quantify these observations regarding the timings of selection, we compared the latency of visual and motor selection time courses across memory-load conditions (load two/four) using a jackknife resampling analysis (Miller et al., 1998; see also: van Ede et al., 2019). As a robust estimate of latency, we used where the time-courses first reached 50% of their peak (i.e., minimum in this case). As expected from looking at the time courses, we found no significant difference in the latency of the visual alpha-band signal between load-two and load-four (**Figure 4A**; *t*(24) = 1.28, *p* = 0.21, *d_z_* = 0.05). In contrast, despite this lack of a difference in the latency of visual selection, we observed a robust latency difference in motor selection, whereby the beta-band lateralization was significantly delayed in load-four compared to load-two (**Figure 4B**; time difference (*M*±*SE*): 145±58 ms; *t*(24) = 2.46, *p* = 0.02, *d_z_* = 0.1).

Together, these data suggests that delayed task reports with higher working-memory load may not be due to it taking longer to orient to and select the appropriate visual representation, but rather to a lower preparedness to act on this representation.

### Variability in the onset of memory-guided behavior covaries with the timing of motor but not visual selection

So far, we have shown that an increase in memory increases decision times (**Figure 2**), and that it delays the selection of motor, but not visual information from working memory (**Figure 4**). Interestingly, the observed delay in motor selection (145±58 ms) also approximates the observed delay in memory-guided behavior (197±24 ms). This raises the question whether and how the timing of visual and motor selection might also co-vary with decision times within load condition. If the main determinator of the speed of memory-guided behavior is our ‘preparedness to act’ rather than ‘item accessibility’, then we should expect to observe that the timing of motor selection, but not visual selection, co-varies with decision times, even *within* each load condition.

To address this, we separated trials in each load condition based on their decision times using a median split, and marked trials as either *fast* or *slow.* This time, rather than comparing load condition, we compared fast and slow trials within each load condition (**Figure 5**, top rows for load-two, bottom rows for load-four), again separately for visual (**Figure 5A**) and motor (**Figure 5B**) selection. We found significant visual and motor selection in all cases (load-two: *fast* cluster *p* = 0.0001, *slow* cluster *p* = 0.0007; load-four: *fast* cluster *p* = 0.0004, *slow* cluster *p* = 0.002).

**Figure 5.**
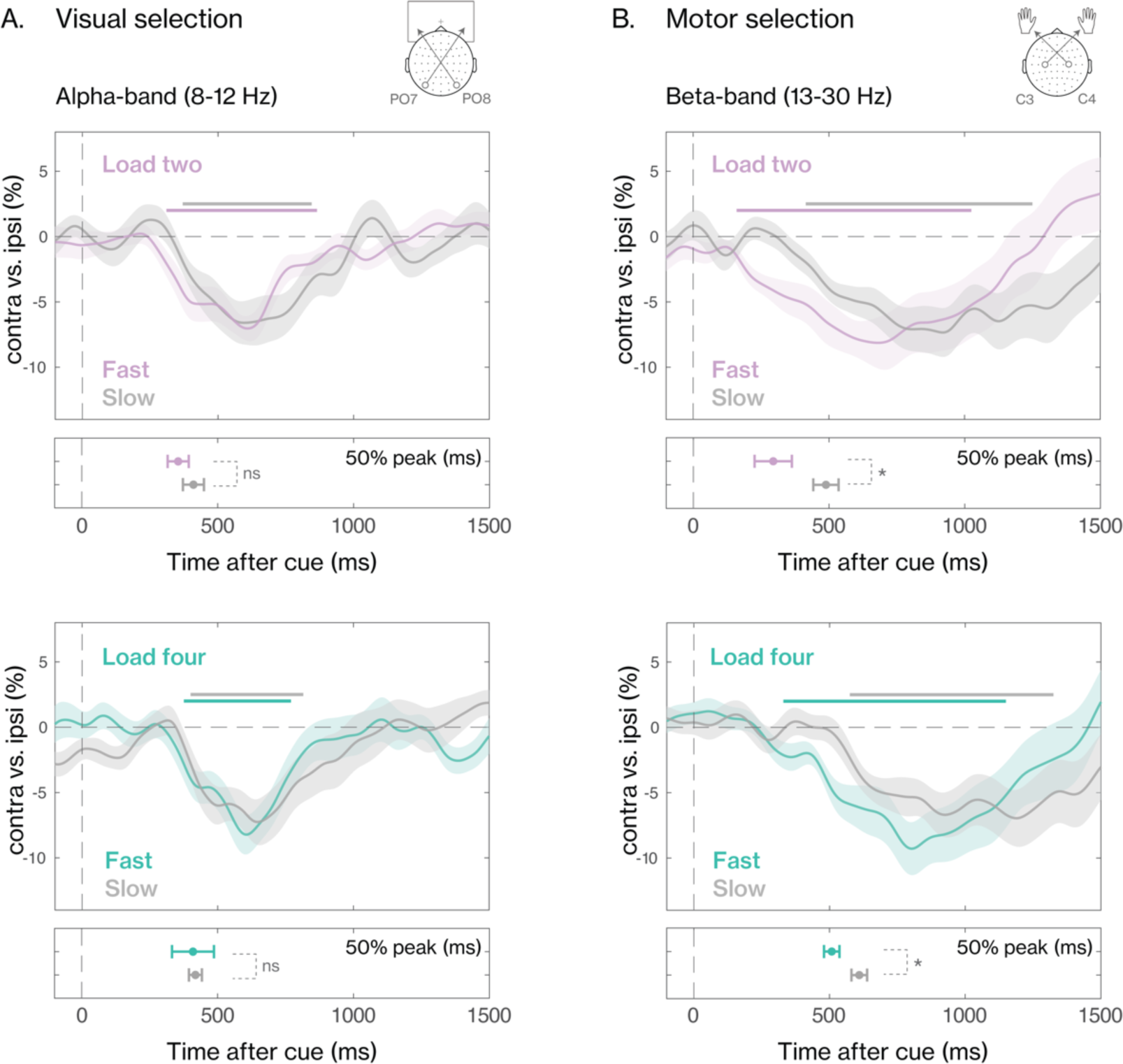
The timing of motor, but not visual selection is delayed in slow compared to fast trials. Time-course of **(A)** the lateralized visual alpha-band (8-12Hz) activity, and **(B)** motor beta-band (13-30Hz) activity, for load-two (upper panel) and load-four (lower panel), and fast (colored) and slow (grey) trials. Thick horizontal lines indicate significant clusters from the cluster-based permutation. Black vertical dashed lines indicate the onset of the fixational color cue. Shadings indicate standard error (SE) across participants. Narrow panels below the time-courses show averaged jackknife estimates of the time the visual and motor time-courses first reach 50% of their minimum (or: negative peak). Error bars indicate standard errors.

Critically, we compared the latencies of visual and motor selection between *fast* and *slow* trials, and again found no significant differences in visual selection (**Figure 5A**), neither in load-two (*t*(24) = 1.38, *p* = 0.2, *d_z_* = 0.05) nor in load-four (*t*(24) = 1.10, *p* = 0.9, *d_z_* = 0.004). In contrast, we again found reliable differences in motor selection (**Figure 5B**), with delayed selection in *slow* compared to *fast* trials, both in load-two (time difference: 194±74 ms; *t*(24) = 2.60, *p* = 0.01, *d_z_* = 0.1) and in load-four (time difference: 101±39 ms; *t*(24) = 2.56, *p* = 0.01, *d_z_* = 0.1).

Together, these data show how the timing of motor selection varies with the onset-speed of memory-guided behavior in both working-memory-load conditions, but the same does not hold for the timing of visual selection. This further supports the idea that the main determinator of the speed of memory-guided behavior is our ‘preparedness to act’ rather than ‘item accessibility’.

## DISCUSSION

It is well established that working-memory load influences task performance – when we hold more items in working memory, we are slower to act upon specific content when it becomes relevant for behavior (e.g., Alvarez & Cavanagh, 2004; Awh et al., 2007; Baddeley, 1992; Bays & Husain, 2008; D’Esposito & Postle, 2015; Luck & Vogel, 1997, 2013; Todd & Marois, 2004; Xu et al., 2018). We asked whether slower responses with higher working-memory load can be accounted for by slower access to sensory representations held in working memory, or by a lower preparedness to act upon them. Here, we show delayed selection of motor information with higher working-memory load (four versus two), but find no evidence for a similar change in the latency of visual selection. Moreover, we show that variability in decision times within each memory load condition is associated with corresponding changes in the latency of motor, but not visual selection. These neural dynamics provide a proof-of-principle that working-memory load can affect our preparedness to act on, without necessarily changing our ability to access, sensory representations in working memory.

It has previously been proposed that a serial internal-comparison process could account for slower responses with higher working-memory load (Sternberg, 1966, 1969, 1975). In this longstanding and prevalent framework, it is posited that contents in working memory are considered serially for selection, such that finding the relevant item takes longer when we hold more items in working memory. While this original line of research considered verbal working memory of letters, it was recently proposed that serial search may also underlie the selection of content in *visual* working memory (Kong & Fougnie, 2019). While these prior studies relied on behavioral measures, we directly tracked the latency of visual and motor selection through lateralized posterior alpha-band and central beta-band activity, respectively. We find that higher working-memory load indeed delays the onset of memory-guided behavior, but that this is met only by a corresponding delay in action selection, without evidence for a similar delay in visual selection. While we do not wish to claim that selection of content in working memory will never be serial, we here find no evidence for the idea that the observed slower responses with higher working memory load can be accounted for by delayed visual selection. Instead, they are better accounted for by delayed selection of the relevant manual action. Accordingly, our results demonstrate that action readiness is another key factor that can account for slower memory-guided behavior with higher working-memory load.

Our conclusions ultimately hinge on the suitability and interpretation of our neural markers for sensory access and action readiness. The lateralization of central beta-band activity is a canonical marker of motor planning and selection (e.g., Mcfarland et al., 2000; Neuper et al., 2006; Salmelin & Hari, 1994; Van Wijk et al., 2009). We interpret the latency of motor selection to reflect the degree to which motor information is readily available. The latency of this signal is not a direct measure of readiness, but should rather be seen as a *consequence*: the higher the degree of action readiness, the faster the relevant manual action can be selected and executed by the participant. Moreover, complementary to the current study, we and others have previously demonstrated that action planning takes place during the working-memory delay alongside visual encoding and retention (Boettcher et al., 2021; Henderson et al., 2022; Rösner et al., 2022; Schneider et al., 2017), predicts future task performance (Boettcher et al., 2021; Nasrawi et al., 2023), and is more pronounced with lower working-memory load (Nasrawi & van Ede, 2022). These previous findings substantiate our inference that the latency of motor selection is likely a consequence of higher action readiness in load two compared to load four.

Likewise, the lateralization of posterior alpha-band activity is a canonical neural marker of visuo-spatial attention (Jensen & Mazaheri, 2010; Liu et al., 2022; Poch et al., 2017; Thut et al., 2006; Worden et al., 2000). While we observed clear alpha lateralization after the cue in both memory-load conditions, it remains unknown whether this signal reflects the process of selecting relevant content from working memory, in addition to spatial orienting towards the cued memory item. Nonetheless, if the selection of content from working memory relied on a mechanism of internal serial search, one should still predict that delayed ‘finding’ of the relevant item with higher working-memory load should concur with delayed spatial orienting to its memorized location – a prediction we find no evidence for.

Evidently, alpha lateralization may not have sufficient temporal resolution to rule out the possibility that serial selection may still occur at a very fast timescale. However, in light of our behavioral decision-time and neural motor-selection delay of approximately 200 milliseconds, such potential small differences in the latency of visual selection unlikely account for the observed delayed responses with higher working-memory load.

In the current study, we focused on the influence of working-memory load on decision times, and additionally compared trials based on decision times within each memory load condition. Another behavioral measure that is affected by working-memory load is task accuracy or precision. Although we also find a lower precision of the orientation-reproduction reports in load four compared to load two, it is unlikely that differences in response readiness also account for this effect. Instead, reduced precision of memory-guided behavior is more likely to be affected by variables such as exceeded capacity (Luck & Vogel, 1997; Todd & Marois, 2004; Vogel & Machizawa, 2004; Zhang & Luck, 2008), reduced resources per item (Bays et al., 2009; Bays & Husain, 2008; Ma et al., 2014; see also: van den Berg et al., 2012), and/or increased inter-item interference (Oberauer et al., 2012; Oberauer & Lin, 2017). While such variable may also contribute to the speed of memory-guided behavior, our study posits response readiness as a key factor that may particularly contribute to this. Moreover, we show that response readiness does not only account for the influence of working-memory load on decision times, but also to trial-by-trial variations in decision times within each memory-load condition. This reinforces the idea that our preparedness to act is the main determinator of the speed of memory-guided behavior, rather than item accessibility.

The current study builds upon and complements a growing body of research which appreciates that working memory is not merely a storage mechanism, but an anticipatory buffer that forms the bridge between perception and action (Chatham & Badre, 2015; Formica et al., 2024; Heuer et al., 2020; Myers et al., 2017; Olivers & Roelfsema, 2020; Rainer et al., 1999; Trentin et al., 2023; van Ede, 2020; van Ede & Nobre, 2023; Vries et al., 2020). We reinforce this action-oriented perspective on visual working memory, and reveal that working-memory load can affect our preparedness to act on, without necessarily changing our ability to access, sensory representations in working memory.

## MATERIALS AND METHODS

### Participants

Twenty-five healthy human adults (age: mean = 23.68, SD = 3.77; gender: 15 female, 10 male; one left-handed) participated in the experiment. All participants had normal, or corrected-to-normal vision. Sample size was determined a-priori to match prior studies that relied on similar outcome measures (Boettcher et al., 2021; Nasrawi et al., 2023; van Ede et al., 2019). No participants were excluded from the analyses. The experiment was approved for by the Research Ethics Committee of the Vrije Universiteit Amsterdam. Participants provided written informed consent before participating in the study, and were rewarded €10 or 10 research credits per hour for their participation.

### Experimental design and procedure

Participants performed a visual-motor working memory task (**Figure 1**) in which participants were asked to memorize the orientation of colored bars, and reproduce the orientation of one of the bars after a delay using a response dial (**Figure 1A**). We build on a recently developed task that was also used in several complementary recent studies (Boettcher et al., 2021; Nasrawi et al., 2023; van Ede et al., 2019), and manipulated two key factors.

First, a blocked working-memory load manipulation was implemented (**Figure 1B**). During each trial, four colored bars were always presented to the participant. However, preceding each block, participants were instructed to selectively encode and memorize either two or four items based on their color (e.g., load-two: orange and purple; load-four: orange, purple, green, blue). This ensured memory load was effectively manipulated, while ensuring equated sensory stimulation across trials. Second, similar to our most recent study (Nasrawi et al., 2023), we manipulated and counterbalanced the tilt-response mapping (**Figure 1C**), and ensured that the bars’ orientation was predictively linked to the response hand needed for reproduction (see details below). Additionally, we ensured that response hand was varied independent of cued item location (left/right). Together, these manipulations enabled us to disambiguate EEG markers of motor selection (i.e., required response hand) from those of visual selection (i.e., location of memorized item) after the cue onset, and to compare these signals across our working-memory load conditions (two/four).

The experiment was programmed in PsychoPy (Peirce et al., 2019). All stimuli were presented on a LED monitor (ASUS ROG STRIX XG248; 23.8 inch, 1920 x 1080 pixels, 240 Hz) situated 60 cm away from the participant. Participants were instructed to keep central fixation, and their fixation was monitored during the task by the experimenter using an eye-tracker (EyeLink 1000 plus). Each trial contained an encoding display, presented for 250 msec, with four colored oriented bars presented at four quadrants of the screen, at 3° visual angle distance from the central fixation cross. Two bars were always tilted to the left, the other two to the right. In load two, the two relevant bars were always positioned opposite and diagonal (left-top and right-bottom, or right-top and left-bottom) to one another, and each had a different tilt direction (left/right). As such, the relevant memory items were always balanced relative to fixation. The magnitude of the orientation of the bars was randomly determined, and varied between 5° and 85° to avoid fully vertical, or fully horizontal orientations. Each bar had a width of 0.4° and height of 3° visual angle, and could have one out of four possible colors: orange (HEX-value: #FF8A65), blue (#64B5F6), green (#81C784), or purple (#CE93D8).

After a working-memory delay (2250 msec), the fixation cross changed color, matching the color of one of the oriented bars. We refer to this as the cue. The color change prompted participants to reproduce the precise orientation of the cued color-matching bar held in memory. Participants were instructed to respond as fast and as precisely as possible. As soon as participants initiated their response, the response dial appeared around the central fixation cross, consisting of a large circle (diameter: 3° visual angle) with two handles placed opposite each other on the circle to represent an orientation (see **Figure 1A**). The response dial could either be turned clockwise or counterclockwise (max 90° in each direction), by pressing and holding the “Z” or “M” key, respectively, using the left and right index finger. Participants could only press one key per trial, and the dial would turn for as long as the participant held the button down. After releasing the key, participants finalized their response, and the final orientation of the response dial was reported. After each trial, participants received feedback on their response precision, which was displayed as the percentage of overlap between the reported and target orientation (0-100%).

Visual tilt was linked to the required response hand throughout the experiment: a leftward or rightward tilted bar could only be accurately reported by pressing and holding the correct key. To achieve this, the starting position of the response dial was fixed during, and varied between, blocks: it could be either vertical, or horizontal as illustrated in **Figure 1C** (but never changed within a block). Participants could only turn the dial handles a maximum of 90 degrees in either direction. As a consequence, when the response dial started in a vertical position, a leftward tilted bar could only be reproduced using the left hand, and a rightward tilted bar using the right hand. Alternatively, when the response dial started in a horizontal position, a leftward tilted bar could only be reproduced using the right hand, and a rightward tilted bar using the left hand. Item tilt was thus always linked directly to the required response hand, but the exact mapping between tilt and hand was varied across blocks. As elaborated further in Nasrawi et al. (2023), this manipulation enabled us to track the selection of motor information (i.e., the required response hand), independent of visual tilt, by collapsing across blocks with the above described two possible tilt-response mappings.

Preceding the main experiment, participants practiced the task for 5–10 min, or until their orientation-reproduction performance reached 75% precision. Participants completed two consecutive sessions of one hour, with a self-paced break in between. Each session consisted of 8 blocks of 64 trials (512 trials per session). Preceding each change of the tilt-response mapping, participants completed four practice trials in which participants practiced turning the response dial clockwise and counterclockwise to match a visible orientation. This was done to minimize confusion about the tilt-response mapping during the main trials. These practice trials were not included in the presented analysis.

### Behavioral analyses

Behavioral data were analyzed using R (R Core Team, 2022). Two behavioral measures were used: absolute error (in degrees) and decision time (in msec). Error was defined as the absolute difference in orientation between the reproduction-report and the cued memory item. Decision time was defined as the time it took participants to initiate their response (i.e., button press), after cue onset. Trials with very fast (<100 msec) of slow (>5000 msec, or > mean + 2.5 time standard deviation) were excluded from the analyses. To evaluate the effect of memory load (two/four) on absolute error and decision times, conventional paired samples t-tests were performed. Data visualization was done using the ggplot2 package (Wickham, 2016). Bar plots and violin plots were used to visualize the effect of load on both behavioral measures. Density plots were used to show the distributions of decision times, and the reported orientation as a function of target orientation, for load-two and load-four.

### EEG acquisition and analyses

#### Acquisition

The BioSemi ActiveTwo System (biosemi.com) was used to measure EEG, with a conventional 10–10 System 64 electrode setup. Two electrodes on the left and right mastoid were used for offline re-referencing of the data. In addition, EOG was measured using one electrode next to, and another above the left eye, making it sensitive to both horizontal and vertical eye-movements.

#### Pre-processing

EEG analyses were performed in MATLAB (2022a; The MathWorks, 2022) using the FieldTrip toolbox (Oostenveld et al., 2011; https://fieldtriptoolbox.org). The data were epoched 1000 msec before, and 3000 msec after, cue onset, re-referenced to an average of the mastoid electrodes, and down-sampled to 200 Hz. Noisy channels were noted down during recording, and interpolated using an average of the two laterally adjacent electrodes. Independent Component Analysis (ICA) was used to clean-up artifacts of blinks and eye-movements in the data. The time-courses of the ICA components were correlated with the EOG time-courses to identify which ICA components should be rejected.

Additionally, the function *ft_rejectvisual* was used to find and remove trials with exceptionally high variance, and trials with very fast/slow decision times (see section: Behavioral analyses) were rejected from further analyses. Trial rejection was done on all trials, without knowledge of the conditions to which the trial belonged. After trial removal, the counterbalancing of trials conditions was restored so that there was still an equal amount trials with a specific visual/motor congruency, based on item location and required response hand. Finally, a surface Laplacian transform (Perrin et al., 1989) was applied (as also done in: Boettcher et al., 2021; Nasrawi et al., 2023; Nasrawi & van Ede, 2022; van Ede et al., 2019) to increase spatial resolution and enhance sensitivity to the motor beta-activity EEG marker.

#### Time-frequency analysis

Time-frequency responses were acquired by subjecting Hanning-tapered data to a short-time Fourier transform. A 300-ms sliding time window was employed to estimate spectral power within the 3 to 40 Hz range (in 1 Hz intervals), in progressive steps of 50 msec.

To obtain our markers of visual and motor selection, we applied equivalent analyses on two aspects of the data. For motor selection, activity in predefined canonical central motor electrodes (C3/C4) was compared between trials where the required response hand was contralateral versus ipsilateral to these electrodes. This comparison was presented as a normalized difference: [(contra-ipsi)/(contra+ipsi)] × 100. The averaged contrasts across the left and right motor electrodes were then computed. To derive time courses of lateralized motor activity, the contralateral versus ipsilateral response within the predefined 13-to 30-Hz frequency band was averaged. Topographies of the lateralized motor activity were generated by contrasting trials associating memory content with a left versus right hand response, expressed as a normalized difference between left and right trials for each electrode. Equivalent to our analysis of lateralized motor selection, we also investigated lateralized signatures of visual selection in standard posterior visual electrodes PO7/PO8, focusing on the predefined 8–12 Hz frequency band. For the analysis of visual selection, we defined left/right and contra-/ipsilateral relative to the memorized location of the cued memory item (as opposed to the associated response hand). As we made sure to independently vary item location and required response hand, the EEG signatures of visual and motor selection were orthogonal by design.

#### Condition and trial-class comparisons

The above described approach was used to calculate visual and motor selection time-courses that we compared between our working-memory load conditions (two/four). In addition, we applied a median split within each memory load condition and participant, by marking trials as fast or slow relative to the median decision time (within this condition and participant). This enabled us to complement our main working-memory load comparison with additional comparisons between fast and slow trials, within each memory-load condition.

#### Statistical evaluation

As in our previous studies, cluster-based permutation analyses (see Maris & Oostenveld, 2007) were employed for the statistical assessment of EEG time–frequency responses (considering clusters in time and frequency) and time-courses (considering clusters in time), utilizing 10,000 permutations and a significance level of 0.025. This non-parametric, Monte Carlo approach, provides a solution for the challenges associated with multiple comparisons in the statistical analysis of EEG data, particularly when dealing with a substantial number of time-frequency comparisons.

In addition, we used a jackknife resampling approach to assess latency differences between EEG time-courses (Kiesel et al., 2008; Miller et al., 1998; Rainer et al., 1999; Ulrich & Miller, 2001; see also: van Ede et al., 2018, 2019). The moment when the time-course signal reached 50% of its maximum (defined as the largest deviation from zero) was estimated iteratively, leaving out one participant at each iteration. We compared the timing of the visual alpha-band and motor beta-band time-courses between memory-load conditions (two/four), and within memory load condition for fast versus slow trials. To statistically evaluate these differences, the standard error of the jackknife-derived difference score is corrected according to: sqrt((nsub-1)/nsub x sum((diff-avg)^2)) (see: Miller et al., 1998), before conventional paired samples t-tests were performed.

## Acknowledgements

This research was supported by an ERC Starting Grant from the European Research Council (MEMTICIPATION, 850636) and an NWO Vidi grant by the Dutch Research Council (grant number 14721) to F.v.E.

## Notes

### Competing Interest Statement

The authors have declared no competing interest.

